# MCT1-dependent energetic failure and neuroinflammation underlie optic nerve degeneration in Wolfram syndrome mice

**DOI:** 10.1101/2022.07.18.500452

**Authors:** Greta Rossi, Gabriele Ordazzo, Niccolò N. Vanni, Valerio Castoldi, Angelo Iannielli, Dario Di Silvestre, Edoardo Bellini, Letizia Bernardo, Serena G. Giannelli, Sharon Muggeo, Leocani Letizia, PierLuigi Mauri, Vania Broccoli

**Affiliations:** Division of Neuroscience, San Raffaele Scientific Institute, 20132 Milan, Italy; Experimental Neurophysiology Unit, Institute of Experimental Neurology (INSPE), San Raffaele Scientific Institute, 20132 Milan, Italy; National Research Council of Italy, Institute of Neuroscience (IN-CNR-IN), 20129 Milan, Italy; National Research Council of Italy, Institute of Technologies in Biomedicine (ITB-CNR), 20090 Milan, Italy

**Author notes:** Corresponding author: Vania Broccoli Stem Cells and Neurogenesis Unit, Division of Neuroscience, San Raffaele Scientific Institute, Via Olgettina 58, 20132 Milan, Italy. Website: www.vaniabroccolilab.com.

## Abstract

Wolfram syndrome 1 (WS1) is a rare genetic disorder caused by mutations in the *WFS1* gene leading to a wide spectrum of clinical dysfunctions, among which blindness, diabetes and neurological deficits are the most prominent. *WFS1* encodes for the endoplasmic reticulum (ER) resident transmembrane protein Wolframin with multiple functions in ER processes. However, the *WFS1*-dependent etiopathology in retinal cells is unknown. Herein, we showed that *Wfs1* mutant mice developed early retinal electrophysiological impairments followed by marked visual loss. Interestingly, axons and myelin disruption in the optic nerve preceded the degeneration of the retinal ganglion cell bodies in the retina. Transcriptomics at pre-degenerative stage revealed the STAT3-dependent activation of proinflammatory glial markers with reduction of the homeostatic and pro-survival factors Glutamine synthetase and BDNF. Furthermore, label-free comparative proteomics identified a significant reduction of the monocarboxylate transport isoform 1 (MCT1) and its partner Basigin that are highly enriched on retinal astrocytes and myelin-forming oligodendrocytes in optic nerve together with Wolframin. Loss of MCT1 caused a failure in lactate transfer from glial to neuronal cell bodies and axons leading to a chronic hypometabolic state. Thus, this bioenergetic impairment is occurring concurrently both in the axonal regions and cell bodies of the retinal ganglion cells, selectively endangering their survival while impacting less on other retinal cells. This metabolic dysfunction occurs months before the frank RGC degeneration suggesting an extended time window for intervening with new therapeutic strategies focused on boosting retinal and optic nerve bioenergetics in WS1.

## INTRODUCTION

Wolfram syndrome 1 (WS1) is a rare and multisystemic genetic disease. In most cases the first clinical sign is the development of non-autoimmune diabetes mellitus (DM) during infancy followed by optic atrophy (OA) which leads to progressive visual loss in adolescence progressing relentlessly to blindness in adulthood (Wolfram and Wagener, 1938; Barret et al., 1995). While DM symptoms are clinically treatable with anti-diabetic therapies, therapies to arrest or delay the loss of sight are lacking (Urano et al., 2016). Although concurrent DM and OA are pathognomonic for WS1, patients might develop additional serious neurological illnesses, most commonly hearing loss, ataxia, epilepsy and peripheral neuropathy (Minton et al., 2003; Chaussenot et al., 2011). Serious psychiatric disturbances have been also described in WS1 patients including psychosis, episodes of severe depression, and impulsive/ aggressive behavior (Switf et al. 1990; Switf and Swift, 2005). OA is caused by the selective loss of retinal ganglion cells (RGCs) and their axons in the optic nerve with a pattern of degeneration that mainly interests the central part of the optic tract rather than the periphery and is accompanied by different degrees of demyelination (Zmyslowska et al., 2019; Barboni et al., 2022). Interestingly, neuroimaging studies revealed diffuse alterations in the gray and white matters of the brain with significant loss of volume of the brainstem and microstructure abnormalities and signs of degeneration of myelin (Lugar et al., 2016; Samara et al., 2019). The gene mutated in WS1 is *WFS1* which encodes for a multi-transmembrane protein and resident in the endoplasmic reticulum (ER) named Wolframin (Inoue et al., 1998; Strom et al., 1998). Mounting evidence suggests that *WFS1* gene loss predispose insulin producing pancreatic β-cells ER stress-mediated apoptosis with disruption of cellular calcium homeostasis (Fonseca et al., 2005, 2010; Riggs et al., 2005; Yamanda et al., 2006). In particular, Wolframin is able to increase the Ca^2+^ uptake within the ER, partially through the regulation of the sarco/endoplasmic reticulum Ca^2+^-ATPase SERCA pump (Zatyka *et al*, 2015). As a consequence, Wolframin loss results in the Ca^2+^ leakage from the ER with a simultaneous increase of its cytosolic concentration, activation of the Ca^2+^-dependent protease calpain which, in turn, activates programmed apoptosis (Lu et al., 2014; Takei et al., 2006). Moreover, recent findings have suggested that Wolframin is required for the proper release of both insulin and neurotransmitters from the ER and their delivery to the cell periphery through assisted vesicle trafficking (Wang et al., 2021). Increasing reports have described dysfunctional ER-mitochondria communication caused by Wolframin loss. In fact, *WFS1* deficiency deranges the inositol 1,4,5-trisphosphate receptor (IP3R)-mediated release by destabilizing the NCS1 protein and, thereby, leading to reduced Ca^2+^ uptake by the mitochondria with consequent functional impairments (Schlecker et al., 2006; Cagalinec et al., 2016; Angebault et al., 2018). Moreover, a recent study unveiled an enriched localization of Wolframin at the MAM sites and in WS1 patient fibroblasts, Wolframin deficiency correlated with MAMs loss and, therefore, with defective Ca^2+^ transfer from the ER to the mitochondria and their related dysfunctions (La Morgia et al., 2020). Altogether, these results highlight mitochondrial dysfunctions as a relevant pathogenetic event triggered by Wolframin and leading to cell death. However, most of these findings were obtained using cell lines and patients’ fibroblasts, and, therefore whether these pathogenetic alterations have the same relevance in neural cells and represent the main and only cause of neurodegeneration remain to be further investigated. Different *Wsf1* targeted mutant mouse and rat lines have been generated and showed to recapitulate the main cardinal pathological traits of the WS patients. Between the mouse models, the neurological and behavioral alterations of the *Wfs1*^exon8del^ targeted mice have been extensively characterized showing diabetic glucose intolerance and anxious-like behaviours upon stressful environment exposure (Kato et al., 2008; Luuk et al., 2008, 2009). Visual impairment in these mice is less characterized with only a histopathological study that reported some thinning of the retinal tissue with lower retinal thickness/longitudinal diameter ratio in 4 month old *Wfs1* deficient mice (Waszczykowska et al., 2020). More recently, a mutant rat line was obtained through *Wfs1* exon 5 disruption showing the hallmarks of WS1, with diabetes, optic atrophy, and neurodegeneration (Plaas et al., 2017). The increase of ER stress levels was detected in mutant retina, brainstem and pancreas, coupled with a volumetric decrease in all these districts. These features made it very useful to test the efficacy of pharmacological treatments (Plaas et al., 2017). Prolonged administration of the glucagon like peptide 1 receptor (GLP1-R) agonists have demonstrated beneficial effects in preventing diabetes, excitotoxicity and ER stress in aged rats and delaying disease progression (Toots et al., 2018; Seppa et al., 2019). Very recently the synergistic effects of GLP1-R agonist and the brain-derived neurotrophic factor mimetic 7,8-DHF co-treatment have shown neuroprotective capacity on rats’ visual pathway rescuing visual activity (Seppa et al., 2021). Thus, rodent models of *Wfs1* deficiency offers pivotal systems where to dissect and further validate the pathogenetic basis of this disease. Herein, we employed the *Wfs1*^exon8del^ mutant mice to better characterize the progressive impairments of visual physiology and activity over time and identify novel molecular and biochemical alterations by genome-wide transcriptomics and proteomics on isolated retinal tissues. These targeted analyses reveal neuroinflammation and bioenergetic substrate loss as the plausible causes for the selective vulnerability of RGCs and their degeneration in WS1 mice.

## RESULTS

### Progressive visual impairments in *Wfs1* mutant mice

To determine RGCs functionality and visual signal conduction to the brain in *Wfs1* mutant mice we exploited whole field photopic electroretinogram (pERG), which measures action potentials produced by the retina when it is stimulated by light of adequate intensity and it is the composite of electrical activity from the photoreceptors, inner retina and RGCs. In photopic conditions, the photopic negative response (PhNR) wave represents the RGC spiking activity originating from light-adapted cones and allows the detection of RGC-specific alterations (Gotoh *et al*, 2004; Kinoshita *et al*, 2016). Whilst PhNR amplitudes resulted unaltered, pERG detected a significantly prolonged implicit time, which refers to the interval between the stimulation and the peak of the negative response, in 8 month old *Wfs1* mutant mice (*WT*: 73 ± 6; *Wfs1-KO*: 82 ± 5; *p* < 0.05) (**Figure 1A**). This early indication of RGC abnormalities became even more pronounced in 12 month old mutant mice in which, beside a delayed response (*WT*: 75 ± 7; *Wfs1-KO*: 88 ± 6; *p* < 0.001) a significant PhNR amplitude decrease appeared evident (*WT*: 10 ± 3; *Wfs1-KO*: 6 ± 2, *p* < 0.001) (**Figure 1A**). To further deepen our neurophysiological evaluation, we sought to investigate light-induced electric conduction from the retina to the brain visual cortex by flash visual evoked potential (fVEP) recordings. These measurements represent a direct readout of the optic nerve functional activity by providing two important parameters: the latency, which measures the propagation speed of the visual stimulus along the nerve fibers until its central detection; the amplitude, that represents the potential difference generated after administration of the visual stimulus, allowing quantification of the power of the electrical signal produced in the optic nerve. VEP trace typically consists of three waves, a first negative wave (N1), followed by both a positive (P1) and a second negative wave (N2), with the first two related to RGC function, whereas the latter to the Müller cell activity (**Figure 1B**). Consistent with our ERG findings, 8-month-old mutant retinas did not present PEV amplitude variations (*WT*: 44 ± 8; *Wfs1-KO*: 42 ± 6) but displayed increased latencies (*WT*: 46 ± 4; *Wfs1-KO*: 52 ± 2, *p* < 0.001), indicative of slower transmissions along the visual pathway (**Figures 1B**). However, both PEV parameters were significantly affected as mouse age progressed (PEV amplitude *WT*: 40 ± 4; *Wfs1-KO*: 28 ± 2; *p* < 0.001; PEV latencies *WT*: 47 ± 2; *Wfs1-KO*: 52 ± 3, *p* < 0.001) (**Figures 1B**). We then asked if these neurophysiological deficits correlate with manifested loss of visual acuity as recorded by the optomotor reflex response (OMR) based on tracking the unconditioned compensatory head movements of the animal while trying to stabilize the image of the moving environment (rotating vertical stripes). *Wfs1* mutant mice showed a progressive reduction in visual acuity with a significant loss already at 8 months which markedly worsened 4 months later (4 months - *WT*: 0.40 ± 0.5; *Wfs1-KO* 0.36 ± 0.8 c/deg, 8 months - *WT*: 0.39 ± 0.4; *Wfs1-KO*: 0.30 ± 0.3 c/deg, *p* < 0.05, 12 months - *WT*: 0.38 ± 0.6; *Wfs1-KO*: 0.19 ± 0.5 c/deg, *p* < 0.01) (**Figure 1C**). These results imply that the initial visual impairment in *Wfs1* mutant mice is represented by a delay of the visual stimulus along the nerve fibers suggesting that optic nerve damage precedes the dysfunctions in the RGC somata. Subsequently, RGC functions are extensively altered causing a drastic loss of visual acuity at 12 months of age.

**Figure 1:**
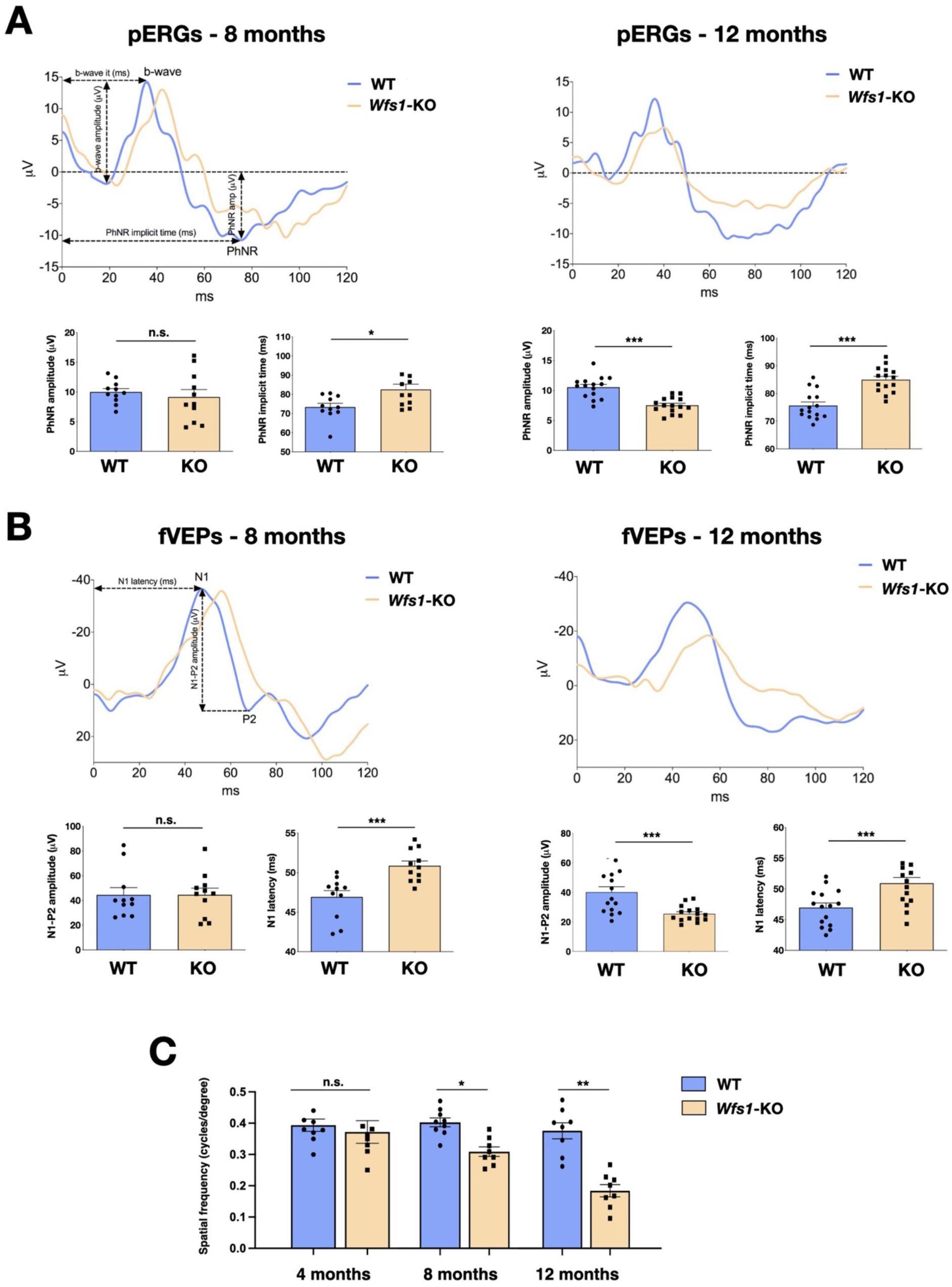
*Wfs1* mutant mice show progressive and severe impairment of visual activity. (A) Representative pERG waveforms at 8 and 12 months of age in wild-type (WT) and *Wfs1* mutant (KO) animals and relative PhNR amplitude and implicit time quantification (n = 15, \**p* < 0.05; \*\**p* < 0.01; \*\*\**p* < 0.001, Student *t*-test). Data are presented as mean ± SEM. (B) Representative fVEP waveforms at 8 and 12 months of age in wild type and *Wfs1* mutant (KO) animals and relative N1 latency and N1-P2 amplitude quantification (n = 15, \**p* < 0.05; \*\**p* < 0.01; \*\*\**p* < 0.001, Welch’s *t*-test for N1-P2 amplitude and Student *t*-test for N1 latency). Data are presented as mean ± SEM. (C) Quantification of visual acuity measuring the opto-motor reflex expressed as cycles per degree in 4, 8 and 12 month old wild-type (WT) and *Wfs1* mutant (KO) mice (n = 8, \**p* < 0.05; \*\**p* < 0.01; \*\*\**p* < 0.001). Data are presented as mean ± SEM.

### Optic nerve degeneration precedes RGC loss in *Wfs1* mutant mice

No deep characterization of the *Wfs1* mutant visual structures has been yet reported and the data available refers exclusively to eye morphology in relatively young animals (Waszczykowska et al., 2020). At first, we evaluated the gross retinal architecture in 8 months old mice by optical coherence tomography (OCT) and fluorescein angiography (FA), in terms of both morphology and neovascularization. FA assessment was necessary to exclude confounding effects due to diabetic retinopathy (DM), which represents one of the major eye complications of DM but whose causative mechanisms are highly different from those in WS1 (Hilson et al., 2009). The collected images did not reveal any signs of retinal vasculature dysfunction in either group of animals. Indeed, OCT-angiography did not highlight any significant difference in the eye fundi or perfusion and vessel length density and tortuosity (**Figure S1**). Moreover, we also assessed the thickness of retinal nerve fiber layer (RNFL), and no significant differences were disclosed by structural OCT (**Figure S1**). Since mutant elderly mice often displayed corneal haze, with consequent no good quality images, we managed to sample only three 12 month old clear-lensed eyes that still did not show big abnormalities in RNFL thickness at OCT segmentation (**Figure S1**). To investigate the cellular dysfunctions underlying the neurophysiological recordings we performed transmission electron microscopy analyses (TEM) of transversal optic nerve sections of *Wfs1* mutant and littermate control mice at the same timepoints (8 and 12 months). Post-hoc TEM imaging analysis enabled us to estimate average RGC axonal densities (mean axon count/area), average myelin area/field also quantifying the empty space (i.e., uncovered by axons) and g-ratio (axonal area divided by axonal area including that of myelin sheets). These elaborations revealed that 8 month old *Wfs1* mutant optic nerves presented incipient myelin thinning (g-ratio - *WT*: 0.52 ± 0.18; *Wfs1-KO*: 0.54 ± 0.14, *p* < 0.05) with reduced myelin area and a higher proportion of empty space, but without axonal count discrepancies (**Figure 2A**). TEM imaging in elder mutant animals (12 months of age) unveiled signs of more severe myelin degeneration (g-ratio - *WT*: 0,42 ± 0,18; *Wfs1-KO*: 0,50 ± 0,18, *p* < 0.05) with pathological features and degenerating fibers (**Figure 2B**). In particular, we observed a significant reduction of axonal density and area, area occupied by myelin together with evident signs of axonal damage with increased g-ratio (*WT*: 0,41 ± 0,04; *Wfs1-KO*: 0,51 ± 0,003, *p* < 0.05) (**Figure 2B**). These results provide evidence that myelin alterations and axonal thinning are the first pathological signs followed by manifested myelin degeneration and axonal loss in *Wfs1* mutant optic nerves. This pathological progression is perfectly matching the timeline of the visual functional deficits previously uncovered by electrophysiological recordings and the optomotor analysis. To assess whether optic axonal damage was occurring with simultaneous alterations in RGC cell bodies within the retinal tissue, we performed a detailed count of RGC number by immunocytochemical staining using the pan-RGCs marker RBMPS (**Figure 2C**). Unbiased stereology counting did not reveal significant RGC number changes in 8 month old *Wfs1* mutant mice (**Figure 2C**). However, 4 months later the loss of the RGC population became evident and statistically relevant (28% ± 6% loss in *Wfs1*-KO vs WT retinas) indicating a progressive loss of RGC somata over time (**Figure 2C**). These results illustrate that axonal damage and myelin degeneration anticipate the frank loss of RGC cell bodies in the retinas, even if within a relative delayed timeframe a severe loss of RGCs can be also detectable in the *Wfs1* deficient retinas.

**Figure 2:**
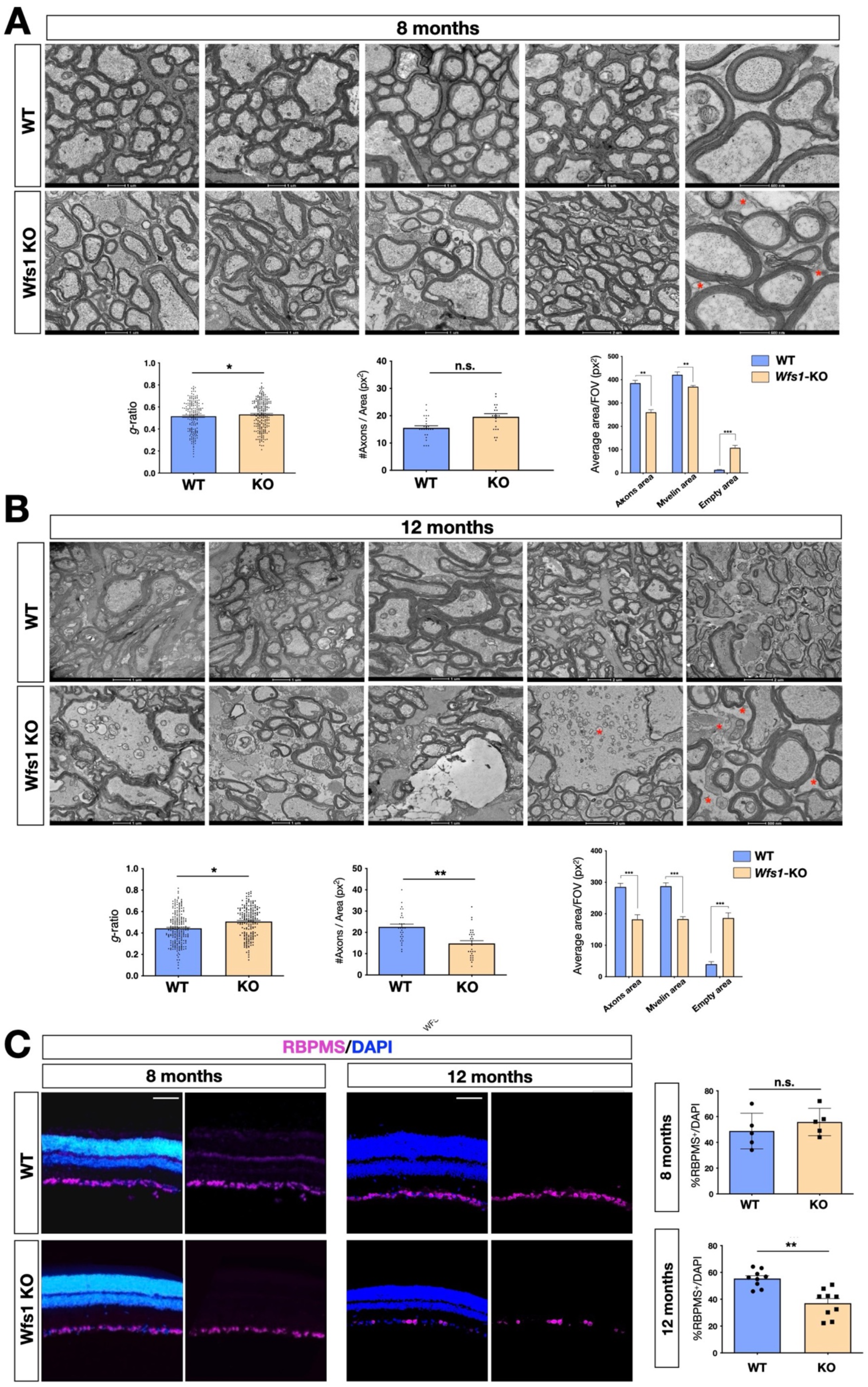
Progressive demyelination in the *Wfs1* mutant optic nerves. (A) Representative EM images of optic nerve cross-sections of wild-type (WT) and Wfs1 mutant mice at 8 months of age showing increased space among the axons (red asterisks). Quantifications of the g-ratio (n = 193, *p < 0.05, unpaired t-test), total number of axons (n = 34), myelin (n = 29) and empty areas (n = 29) normalized by total area (n = 29) (\*\**p* < 0.01; \*\*\**p* < 0.001, 2-way ANOVA with Bonferroni’s post-hoc test). Data are presented as mean ± SEM. (B) Representative EM images of optic nerve cross-sections of wild-type (WT) and Wfs1 mutant mice at 12 months of age showing dramatic expansion of space among the axons (red asterisks). Quantifications of the g-ratio (n = 192, *p < 0.05, unpaired t-test), total number of axons (n = 29), myelin (n = 27) and empty areas (n = 27) normalized by total area (n = 27) (\*\*\**p* < 0.001, 2-way ANOVA with Bonferroni’s post-hoc test). Data are presented as mean ± SEM. (C) Representative immunohistochemical images of retinal cross-sections of wild-type (WT) and Wfs1 mutant (KO) animals at 8 and 12 months of age with RBPMS and DAPI staining in purple and blue, respectively. Relative quantification of RBPMS positive cells over total cell number in the GCL at 8 months (n = 9, *p* = 0,39, 2-tails unpaired t-test) and at 12 months (n = 8, \*\**p* < 0.01, 2-tails unpaired *t*-test. Scale bar, 50 µm.

### Transcriptomic analysis reveals aberrant gliosis which precedes neuronal loss in *Wfs1* mutant retinas

To unveil specific molecular alterations underlying the pathological deficits caused by *Wfs1* gene loss, we performed transcriptome analysis of isolated mutant and control mouse retinas. To investigate prodromic events that anticipate the morphological changes, we performed RNA-seq analysis on tissues from 8 month old mice. RNAs were extracted from whole retinas of three different mice per group (3 WT vs 3 *Wfs1*-KO) and subjected to global gene expression analysis. t-SNE analysis of the RNA-seq datasets confirmed the expected closer similarity and correlation between all mutant retinas in comparison to the analogous wild-type samples (**Figure 3A**). Computational analysis identified 396 genes differentially expressed (DEGs) in mutant retinas compared to the control counterpart, with a similar proportion of either up- or downregulated genes (**Figure 3B**), suggesting not profound transcriptome alterations but rather selective changes in mutant samples at this stage. Interestingly, gene enrichment analysis (Gene Ontology, GO) revealed many deregulated genes related to pathways like cell death regulation, UPR response and protein folding (**Figure 3C**). Intriguingly, all these pathways were down-regulated in the mutant retinas, in line with the general assumption from literature that Wolframin loss impairs the cellular stress-coping capacities. On the contrary, mutant retinas upregulated key genes of neuroinflammatory pathways (**Figure 3C**) as those associated with reactive astrocytes like, Gfap, Lcn2, Edn2, Alpk1 and C4 (**Figure 3D**) (Liu et al., 2022; Linnerbauer et al., 2022). We, next, performed immunocytochemical analysis confirming the aberrant activation of the ER stress effectors ATF6 (*WT*: 0,5 ± 0,3; *Wfs1*-*KO*: 1,5 ± 0,2) and CHOP (*WT*: 0,2 ± 0,01; *Wfs1*-*KO*: 0,4 ± 0,02) in mutant retinal samples (**Figure 4A**). Moreover, GFAP (8 month *WT*: 0,5 ± 0,2; *Wfs1*-*KO*: 1,4 ± 0,3; 12 months *WT*: 0,2 ± 0,02; *Wfs1*-*KO*: 52 ± 2) and VIMENTIN (VIM) (8 months *WT*: 20 ± 2; *Wfs1*-*KO*: 38 ± 5; 12 months *WT*: 2,2 ± 0,5; *Wfs1*-*KO*: 4,6 ± 0,8) staining was found strongly upregulated in mutant retinal Müller glia at both 8 and 12 months of age (**Figure 4B**). Furthermore, the astrocyte-specific Glutamine synthetase (GS) enzyme was found downregulated both at mRNA and protein levels in *Wfs1* deficient retinas (**Figures 3C,4D**). GS generates glutamine by assembling glutamate and ammonia and, therefore, having a critical role in preventing glutamate-dependent excitotoxicity and ammonia toxicity (Rose et al. 2013). Thus, GS reduction in *Wfs1* deficient astrocytes might favor a toxic environment detrimental for neuronal survival. Next, the genes encoding for the key neurotrophic molecule BDNF, and the survival factors Nr4a1 and Nr4a3 were also reduced in *Wfs1* mutant retinas (**Figure 3D**), further altering neuronal homeostasis and stress resilience. In fact, BDNF is a well-known neurotrophic factor expressed by RGCs and muller glia in adulthood retina, where it exerts its autocrine or paracrine effects. Multiple lines of evidence suggest that BDNF promotes survival of RGCs and can ameliorates RGC death after traumatic optic nerve injury (Bennett *et al*, 1999). It is thus not surprising that its downregulation occurs before ganglion cell death. Finally, a number of microglial pro-inflammatory genes such as Il-18, Gpr37, Cybb, Lyz2, Lglas3 and Mpp12 were up-regulated, suggesting that *Wfs1* mutant microglia activate at least some reactive specific pathways (**Figure 3D,E**). However, levels of interferon-inducible genes and NF-κB signaling components were unchanged suggesting the lack of a generalized inflammatory response, but rather a selective activation of specific neuro-inflammatory pathways. To further strengthen our results, RT-qPCR analysis were performed on independent biological samples and substantially confirmed the transcriptome data for the key selected genes (**Figure 3E**), further highlighting a selective pro-inflammatory profile in *Wfs1* deficient retinas. On this line, we noted that the dysregulated Lcn2 and Vim genes in Wsf1 mutant samples have been shown to be direct targets of the STAT3 signaling (Herrmann et al., 2008). Intriguingly, previous studies reported that STAT3 can be stimulated by the ER stress-dependent Perk and Ire1alpha pathways in astrocytes and cancer cells (Meares et al., 2014; Chen and Zhang, 2017). Thus, we profiled by Western blotting both total and active (pY705) forms of STAT3 in control and *Wfs1* mutant retinal lysates at 8 months of age. Interestingly, the active STAT3 variant was highly enhanced in mutant samples although with some variability among the samples (**Figures 5A**). Thus, we profiled by candidate gene expression other STAT3 regulated genes such as FGF2, Il-6, VEGFa, C1s and C1rl finding them consistently up-regulated in *Wfs1* mutant retinas (**Figure 5B**). Altogether, these findings uncovered a prominent pro-inflammatory molecular program associated with reduction of survival and homeostatic factors which is promoted at least in part by the ER stress-dependent STAT3 activation.

**Figure 3:**
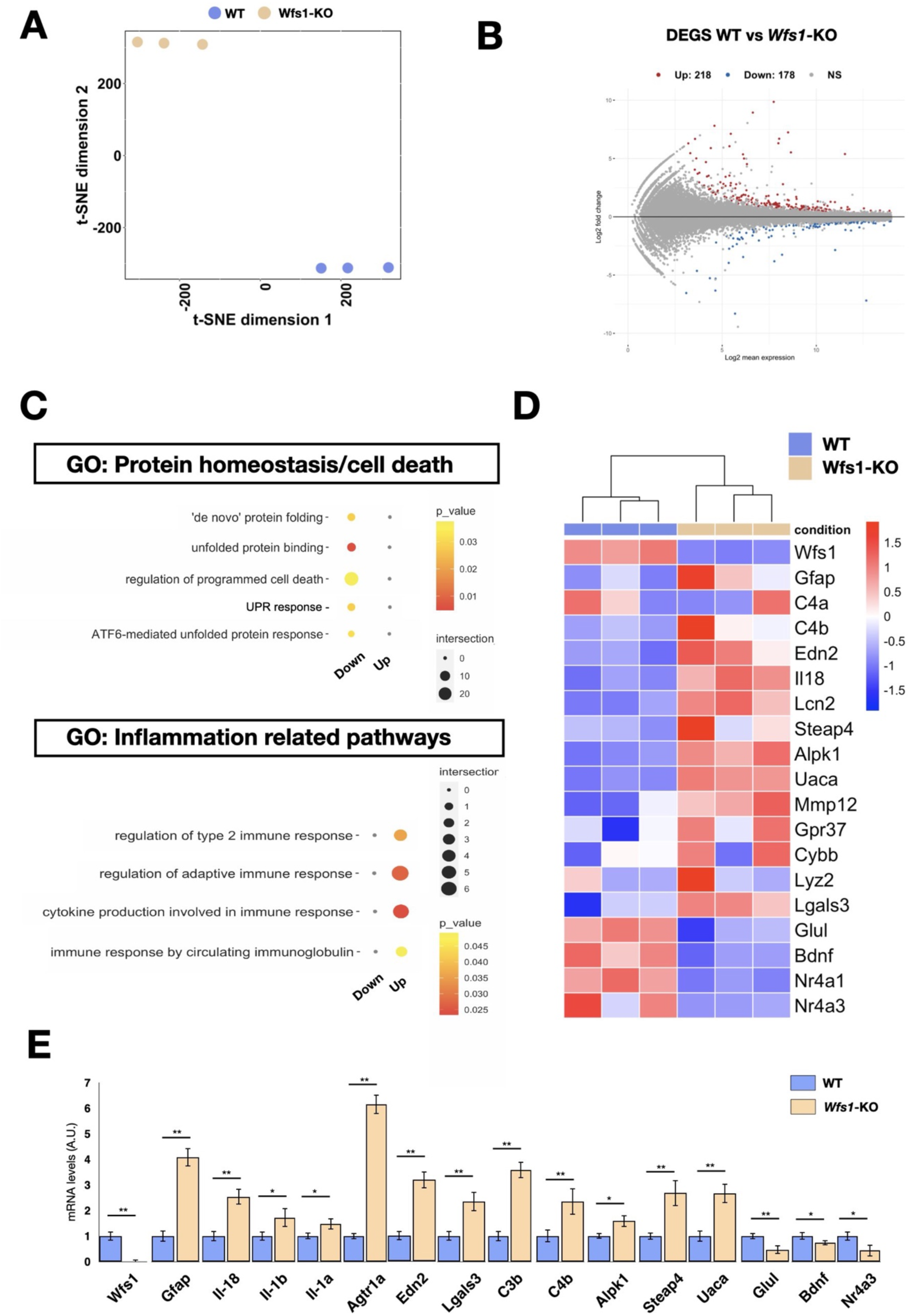
Transcriptome analysis of Wfs1 mutant and control retinas. (A) Whole-transcriptome analysis using the t-Distributed Stochastic Neighbor Embedding (t-SNE) analysis with a view of the sample distribution along the first two dimensions. (B) Significantly up-(red) and down-(blue) regulated genes among the transcriptomic profiles of the samples, shown as highlighted dots in the MA plots. Number of differentially expressed genes (DEGs) (up = 218; down = 178). (C) GO-term categories relative to transcriptional analysis, as in RNA-seq dataset calculated in the list of genes downregulated in WFS1 mutant retinas. (D) Heatmap showing genes normalized count RPKM associated with inflammatory pathways. (E) Boxplot depicting the distribution of gene expression levels, confirming their trends in RNAseq. (n = 3 \**p* < 0.05; \*\**p* < 0.01; \*\*\**p* < 0.001, Student *t*-test).

**Figure 4:**
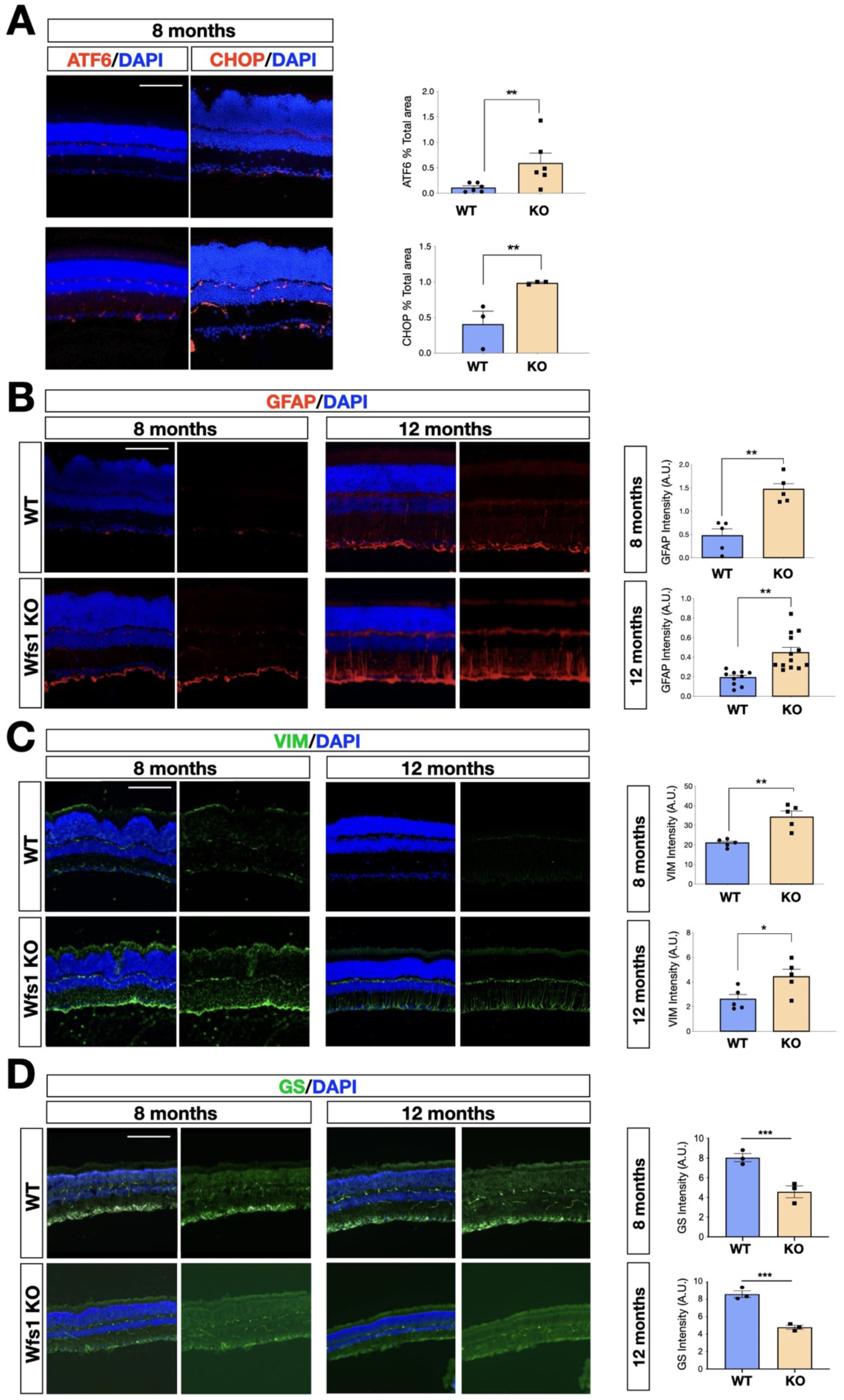
Representative immunohistochemical analysis in wild-type and *Wfs1* mutant retinal tissues at 8 and 12 months of age. (A) Representative images and relative quantification of ATF6 and CHOP immunofluorescence signal in wild-type (WT) and *Wfs1* mutant (KO) retinas at 8 months of age. On the right, quantification of the signal (n = 5, ATF6; n = 12, CHOP, \*\**p* < 0.01, Student *t*-test). Data are presented as mean ± SEM. Scale bar, 100 µm. (B) Representative images of GFAP immunofluorescence in 8 and 12 month old retinas. On the right, quantification of the signal (n = 5, 8 months; n = 13, 12 months, \*\**p* < 0.01, Student *t*-test). Data are presented as mean ± SEM. Scale bar, 100 µm. (C) Representative images of Vimentin (VIM) immunofluorescence in 8 and 12 month old retinas. On the right, quantification of the signal intensity of Vimentin in wild-type (WT) and *Wfs1* mutant (KO) retinas. (n = 5, \**p* < 0,05, \*\**p* < 0.01, Student *t*-test). Data are presented as mean ± SEM. Scale bar, 100 µm. (D) Representative images and relative quantification of the Glutamine Synthetase (GS) immunofluorescence signal in wild-type (WT) and *Wfs1* mutant (KO) retinas at 8 and 12 months. Data are presented as mean ± SEM. (n = 3, \*\**p* < 0.01, Student *t*-test). Scale bar, 100 µm.

**Figure 5:**
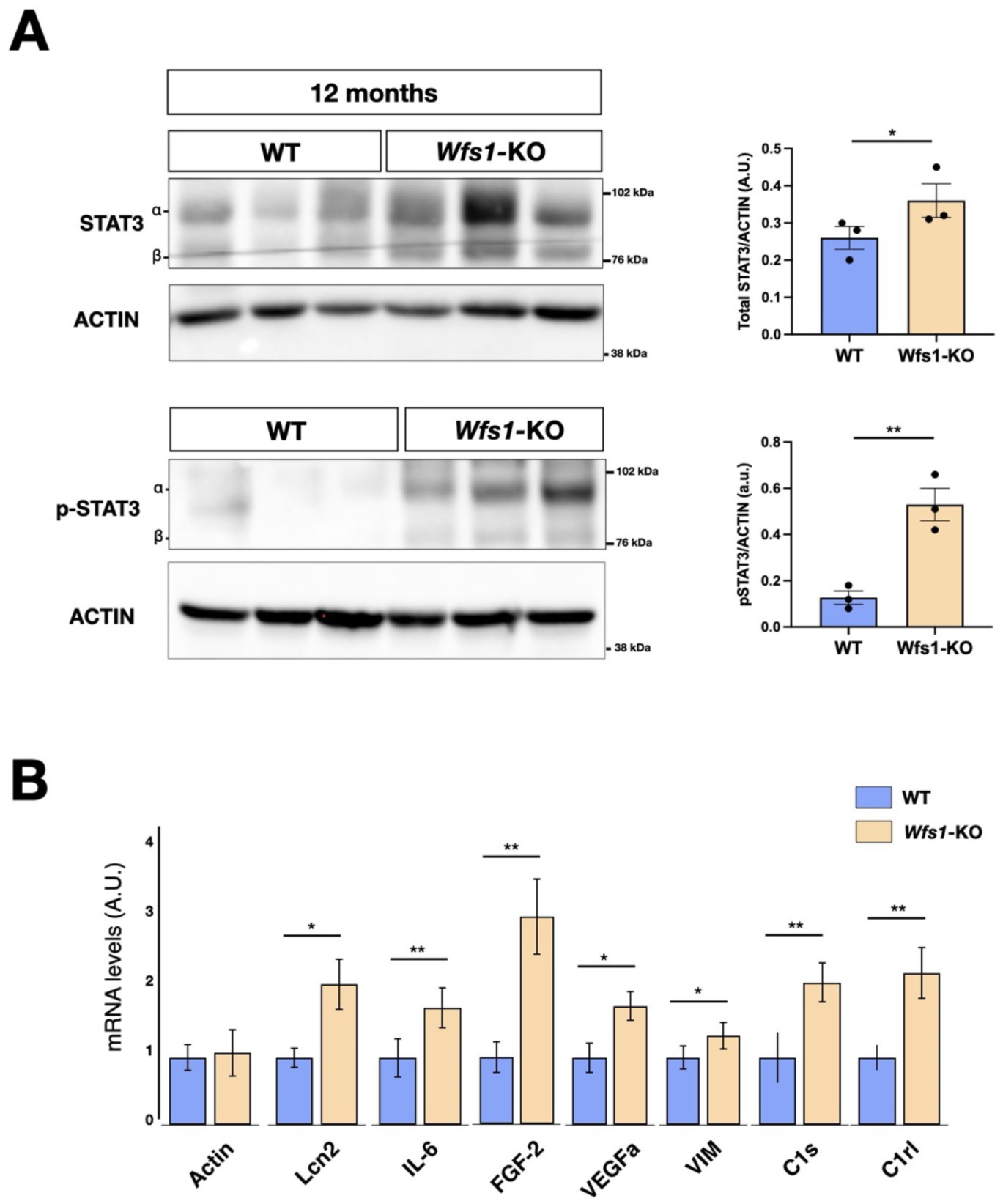
Impaired Signal Transducer and Activator of Transcription 3 (STAT3) signaling in *Wfs1* mutant mice. (A) Representative immunoblotting of pSTAT3 and total STAT3 endogenous protein levels in adult retinae from WT and WFS1 KO mice at 12 months of age. The ratio of p-STAT3 versus total signal transducer and activator of transcription 3 (STAT3) is shown in the graph after normalization over the respective protein load (ACTIN). Quantification of pSTAT3/STAT3 protein signal intensity was performed in 12 months mice retinae (eyes from n = 3 animals, \**p* < 0.05; \*\**p* < 0.01, Student *t*-test). Data are presented as mean ± SEM. (B) Boxplot depicting the distribution of expression levels of STAT3 regulated genes (*p* < 0.05; \*\**p* < 0.01; \*\*\**p* < 0.001, Student *t*-test).

### Proteomics analysis of *Wfs1* mutant retinas

Given that relative few genes showed altered expression levels in *Wfs1* mutant retinas, we postulated that additional key changes can be determined only at protein level. In fact, Wolframin regulates the function of several proteins that control ER physiology and ER-mitochondrial interactions through post-transcriptional mechanisms. Thus, we collected eyes from WT and *Wfs1* mutant mice at 8 months and isolated the retinas with their optic nerves to carry out label-free LC-MS comparative proteomics (**Figure 6A**). Three biological replicates for each of the two genotypes were analysed in triplicate and aligned through Multidimensional Algorithm Protein Mapping (Comunian et al., 2011). Overall, 2046 proteins were identified (**Table S1**) and the virtual 2D map showed the ability of the shotgun approach to identify proteins in a wide range of molecular weight and isoelectrical point (**Figure S2**). In this study, a label-free quantification was applied, considering the total number of peptides (evaluated as Peptide Spectrum Matches or Spectral Count) for each protein. The quantitative analysis, combined with statistical evaluation, led to the identification of 21 stringent differentially expressed proteins between the two genotypes with 9 upregulated and 12 downregulated in the *Wfs1*-KO compared to the WT condition (**Figure 6B** **and Table S1**). Notably, the most upregulated protein upon *Wfs1* gene loss was GFAP, confirming increased gliosis described earlier in the mutant retinas (**Figure 6B**). Conversely, among the most downregulated molecules in mutant samples, we identified MCT1 (Slc16a1) and Basigin. MCT1 belongs to the family of monocarboxylate transporters (MCTs) that promote the transport across cell membranes and the shuttling between cells of energy metabolites as lactate, pyruvate or ketone bodies (Felmlee et al., 2020; Bosshart et al., 2021). Basigin (known also as CD147) is a glycoprotein which assembles with MCT1 promoting its trafficking from ER and its stabilization on the cell membrane (Kirk et al., 2000). In brain, MCTs govern the extracellular release of lactate from glycolytic astrocytes and its transfer into neurons where it is oxidized to generate ATP (Brooks, 2018; Magistretti and Allaman, 2018). This astrocyte-to-neuron lactate shuttle strengthen the metabolic coupling between glial and neuronal cells providing a rapid and tunable energy substrate to sustain the highly dynamic metabolic demand of neurons. To confirm the proteomics results, Western blotting for MCT1 was performed on independent retinal tissue samples confirming a strong downregulation of this transporter both at 5 and 8 months of age (**Figure 6C**). Similarly, a marked reduction of Basigin was detectable in mutant retinas at 5 months of age (**Figure 6C**). Thus, loss of MCT1 and Basigin is detectable months in advance respect to the frank degeneration suggesting that the pathological process proceeds with a relatively slow progression before leading to clinically relevant symptoms. In contrast, MCT2 levels were found upregulated in 8 month old mutant retinas (**Figure 6C**). To determine the causal relationship between Wolframin and MCT1, we transiently expressed MCT1-V5 and Wolframin-mCherry forms in HEK293 and performed co-immunoprecipitation experiments. Interestingly, MCT1-V5 was efficiently retrieved from the Wolframin-mCherry immunoprecipitated samples indicating that the two proteins can associate together in the same protein complex (**Figure 6D**).

**Figure 6:**
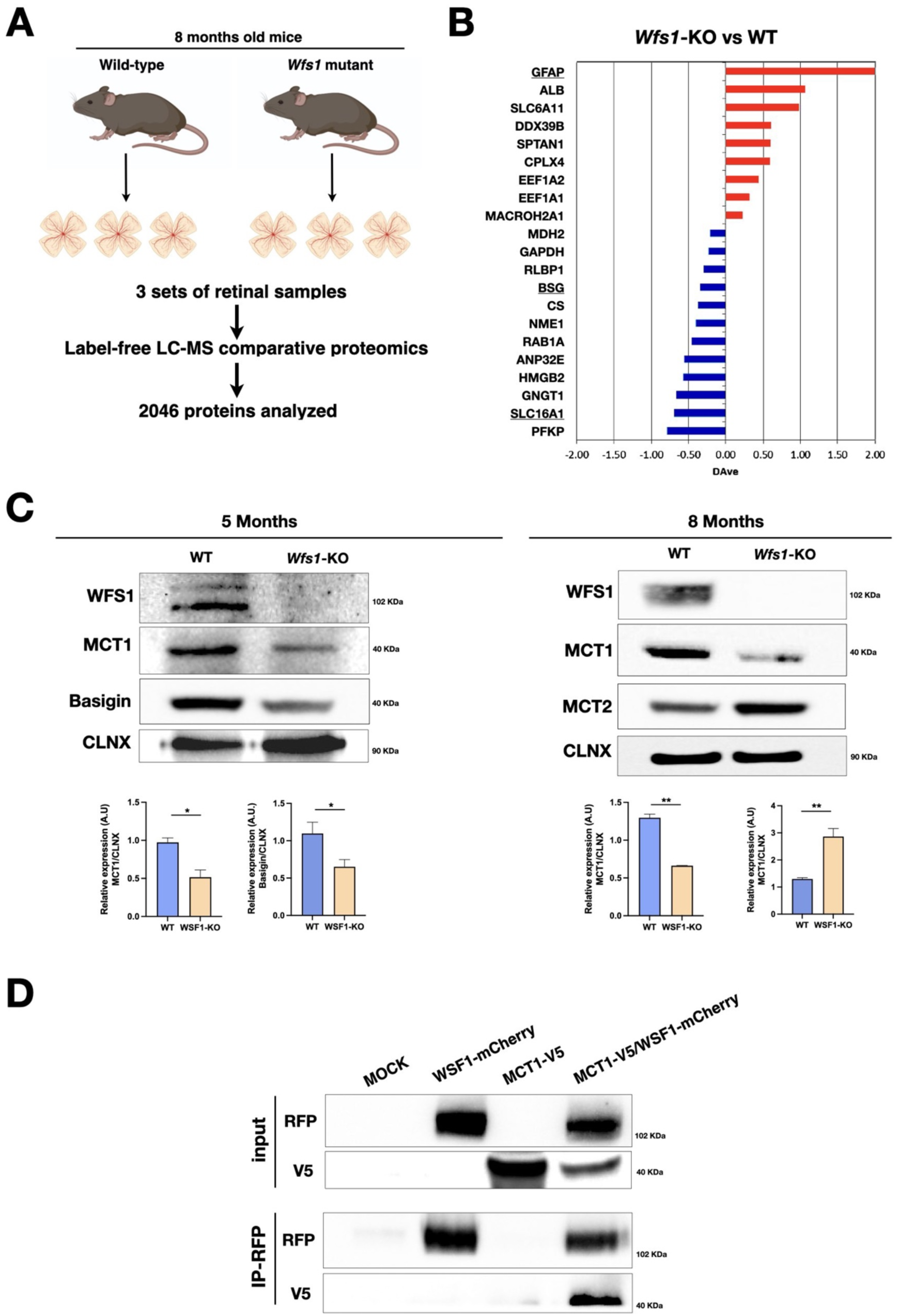
*Wfs1* mutant retinas show altered MCT1, MCT2 and Basigin protein levels. (A) Schematic representation of the label-free LC-MS comparative proteomics approach used to analyze 3 set of retinae tissue samples from WT and Wfs1 mutant mice at 8 months of age. Proteomics analysis of biological replicates for each of biological conditions identified 2046 total proteins. (B) Differentially expressed proteins comparing wild-type (WT) and Wfs1 mutant (KO) mice at 8 months with p-value more significant than 0.05. The positive/red and negative/blue values indicate proteins with levels higher in KO or WT conditions, respectively. (C) Representative western blot images and analysis of Basigin, MCT1 and MCT2 signals in 5- and 8-months mice WT and WSF1-KO retinae with their relative quantification. Calnexin (CLNX) was used as loading control. Data are presented as mean ± SEM (Student t-test, \**p* < 0.05; \*\**p* < 0.01). (D) Representative IP analysis from HEK293T transfected cells with the indicated constructs: WFS1-mCherry and MCT1-V5. RFP-immunoprecipitation was performed, and blot revealed for RFP and V5, showing the proposed interaction between WSF1 and MCT1 in co-transfected condition.

### MCT1 loss causes lactate shuttle defects in *Wfs1* mutant retina and optic nerve

Since each MCT isoform has different preferential localization on neuronal or glial cells and *Wfs1* expression in retina is not precisely defined, we next exploited a recently generated scRNA-seq of adult retinal tissues (Fadl et al., 2020) to determine their expression patterns at single cell resolution. Interestingly, *Wfs1* and *Slc16a1* were found predominantly co-expressed in astroglial and photoreceptor cells (**Figure 7A**). Conversely, expression of *Slc16a7*, encoding for MCT2, was restricted to the retinal neuronal lineage similarly to its selective cell type expression in the brain (**Figure 7A**) (Pierre and Pellerin, 2005). We also confirmed that both Wolframin and MCT1 proteins were detectable in lysates of isolated optic nerves (**Figure 7B**). Immunofluorescence analysis on optic nerve sections confirmed that Wolframin was mainly localized in MBP positive oligodendrocytes wrapping the axonal fibers (**Figure 7C**). Seminal findings have shown that myelin sheaths play a key role in glial-axonal metabolic support by shuttling metabolites like lactate and pyruvate toward axons to fuel the local energy demand (Fünfschilling et al., 2012; Lee et al., 2012). This trophic support is mediated by both MCT1 and MCT2 localized into oligodendroglia and axons, respectively (Halestrap et al., 1999). This process is crucial for energy homeostasis since MCT1 specific inactivation in oligodendrocytes or Schwann cells leads to late-onset axonal degeneration (Jha et al., 2020; Philips et al., 2021). Thus, it is plausible that *Wfs1* gene loss might affect MCT1 protein levels in the optic nerve myelin and the transfer of lactate within the axonal tracts. If this is the case, it is expected that lactate should accumulate in oligoglial cells that are not able to metabolize themselves the large intracellular quantities. Biochemical quantifications in both retinas and isolated optic nerves showed a prominent accumulation of lactate in *Wfs1* mutant samples (**Figure 7D**). Given the failure to traffic lactate outside the oligoglial cells, both *Wfs1*-deficient cell bodies and axons of RGCs are concurrently deprived of a key energy metabolite. In fact, ATP levels were found significantly downregulated in *Wfs1* mutant optic nerve and retinal lysates (**Figure 7E**). Thus, these findings unveil a key role of Wolframin in maintaining physiological levels of MCT1 on glial cells to support the correct transfer of energy metabolites both in cell bodies and axons of RGCs (**Figure 7F**).

**Figure 7:**
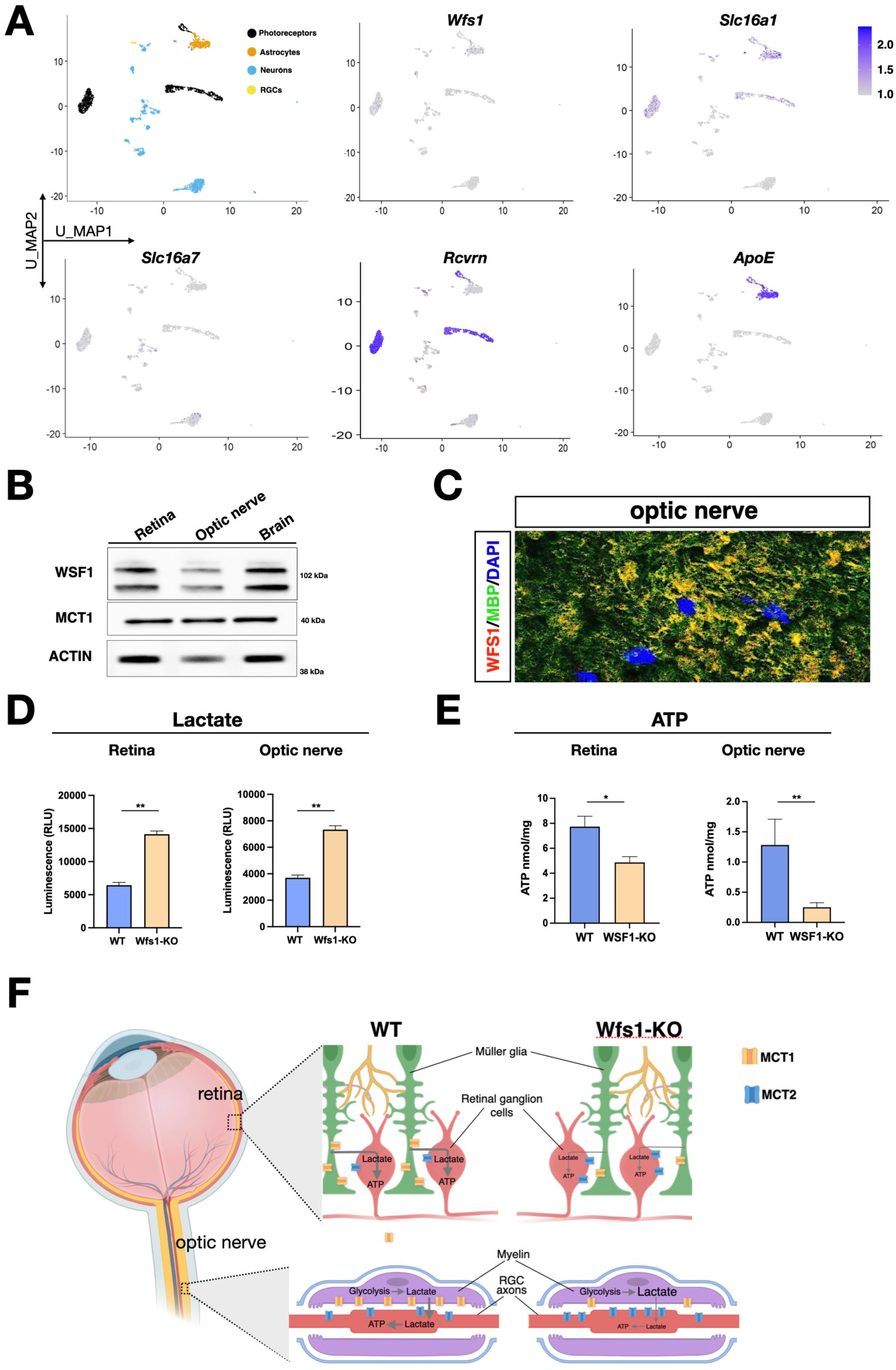
Metabolic alterations in *Wfs1* mutant retinas and optic nerves caused by MCT1 downregulation. (A) Umap of the GSE153673 dataset of scRNA-seq of wild-type (WT) mouse retina showing in the first panel the main cell type composition colored by cluster followed by feature plots colored by expression level of the genes of interest (log2 expression). (B) Representative Western blot showing Wolframin and MCT1 protein levels in wild-type (WT) murine retinal, optic nerve and brain lysates. ACTIN was used as loading control. (C) Representative image of colocalization of WFS1 (red) and MBP1 (green) in WT mice optic nerve section. Scale bar, 10µm. (D, E) Quantification of lactate (D) and ATP (E) in WT and *Wfs1* KO mice retinae and optic nerve shows an increase of lactate level and a reduction of ATP level in *Wfs1* KO mice, respectively. Data are presented as mean ± SEM. (\**p* < 0.05; \*\**p* < 0.01, Student *t*-test). (F) Illustration of the key role of Wolframin in maintaining physiological levels of MCT1 on glial cells to support the correct transfer of energy metabolites in WT retina and optic nerve. In *Wfs1* mutant retina and optic nerve, MCT1 loss results in a concomitant accumulation of lactate and loss of ATP with compensatory increased levels of MCT2 on neuronal cell bodies and axons.

## DISCUSSION

Herein, we exploited unbiased transcriptomics and proteomics quantitative profiling coupled with biochemical and molecular analyses on isolated retinal tissues to disclose new mechanistic insights on RGC degeneration occurring in the WS1 mouse model. In fact, although ER stress and mitochondrial dysfunctions are unequivocally associated with *Wfs1* inactivation, they alone cannot be the cause of the specific cell type loss within the retina occurring in this disease. The retina is a complex assembly of multiple neuronal cell types and photoreceptors that are expected to be comparably sensitive to ER and mitochondrial functional impairments. Herein, we provided unprecedented findings revealing the exact molecular alterations caused by the chronic neuroinflammatory state and the pervasive bioenergetic failure occurring before the development of severe visual deficits in *Wfs1* mutant mice. Beyond the upregulation of prototype markers of inflammation in astrocytes and microglia, we also showed the downregulation of GS and BDNF that are key molecules in glial homeostasis and cell survival. GS catalyzes the conversion of ammonia and glutamate to glutamine, a precursor of glutamate and GABA, which shuttles from astrocytes to neurons (Jayakumar and Norenberg, 2016; Rose et al., 2013). Thus, GS impairment might lead to reduce clearance of extracellular brain glutamate and ammonia, glutamine deficiency, and perturbed glutamatergic and GABAergic neurotransmission. Although these findings identify a new pathological mechanism in WS, this impairment is expected to equally affects retinal cells and to lie rather downstream to the chain of pathological events. Unbiased proteomics profiling of retinal and optic tissue samples enabled us to identify MCT1 and Basigin as significantly downregulated upon *Wfs1* inactivation. MCT1 is the main monocarboxylate transporter in glial cells that promote the shuttling of lactate toward neurons to sustain their elevated energy demand. Intriguingly, we confirmed a simultaneous upregulation of MCT2, the isoform predominantly localized on neuronal cell bodies and axons indicating a compensatory response of the target cells to maximize the entrance of lactate despite its poor extracellular availability (**Figure 7F**). MCT1 loss was detected both in *Wfs1* mutant retinal and optic nerve samples with concomitant accumulation of lactate and loss of ATP. Notably, we showed that *Wfs1* expression is enriched in retinal astrocytes and Wolframin co-localizes with oligodendrocyte-forming myelin in the optic nerve confirming the co-expression with MCT1 at single-cell resolution. Among all retinal neurons, RGCs are the unique neuronal cells with myelinated axons that travel long distances within the optic nerve to reach their brain targets. Thus, these neurons are predicted to have much higher metabolic needs respect to retinal interneurons that feature short-distance and unmyelinated projections contained within the retina. Thus, we postulate that the simultaneously loss of MCT1-dependent lactate transfer from glial cells to cell bodies and axons of RGCs represent a key pathological event which predominantly undermine the survival of this specific retinal cell type in *Wfs1* mutant animals. On this view, mice with only one *MCT1* gene copy develop axonal degeneration by 8 months of age and selective MCT1 inactivation in oligodendrocytes causes axonal injury (Lee et al., 2012; Philips et al., 2013). Thus, these results in mouse together with observations of MCT1 expression alterations in human neurodegenerative disease (Lee et al., 2012; Andres Benito et al., 2018; Tang et al., 2019), strongly point to MCT1-dependent bioenergetic loss as causative for neuronal function alterations and neurodegeneration. Although we described metabolic alterations both in retina and optic nerve in *Wfs1* mutant animals, the first detectable pathological signs are the delay in electric conductance and myelin derangement in the optic nerve. These observations indicate that the optic nerve damage and axonal functional impairment likely represent the initial pathological defects caused by *Wfs1* deficiency. Remarkably, Barboni and colleagues (2022) have recently described that in WS1 patients the RNFL thickness shows a fast decline since early age which precedes of about a decade the atrophy of the cellular bodies of RGCs. Thus, experimental findings in mice and patients converge in highlighting RGC axonal sufferance as the first disease sign in WS1. Moreover, our results uncover chronic bioenergetic failure as an unprecedented new pathological mechanism at the basis of the selective loss of RGC responsible for visual loss in WS1.

Neuroinflammation and MCT1-dependent energy insufficiency represent likely two independent phenomena triggered by separate *Wfs1* dependent pathways. However, chronic inflammation causes a loss of astrocyte-oligodendrocyte gap junctions (Markoullis et al., 2014). These inter-cellular gap junctions enable the shuttle of glucose from the blood circulation to oligodendrocytes through astrocyte intermediates (Wasseff and Scherer, 2011). Thus, astrocytosis can reduce glucose availability in oligodendrocytes and consequently its metabolic support to axons. On this view, MCT1 downregulation and astrocytosis create a pathological axis that synergistically impairs oligodendrocyte metabolic support capabilities.

Several hypotheses can provide an explanation of the loss of MCT1 and Basigin based on the known functions of Wolframin. First, Wolframin was shown to direct vesicle trafficking from ER to the cell periphery and this function can sustain the delivery of specific client proteins (i.e. MCT1 and Basigin) to the cell membrane. Second, Wolframin can stabilize MCT1 and Basigin on the membrane and therefore protect them from selective degradation. Finally, Wolframin-dependent ER stress might particularly affect the stability of selective proteins. An additional hypothesis is that Wolframin has a general facilitatory role in the assembly of oligomeric protein assemblies. In fact, a previous study reported the concomitant loss of both alpha and beta-subunits of the Na^+^/K^+^ ATPase suggesting a possible role of Wolframin in the correct folding and assembly of multimeric complexes in ER (Zatyka et al., 2008). Future studies will be necessary to determine if one or more of these mechanisms are responsible for MCT1/Basigin protein reduction upon Wolframin loss. More in general, it is plausible that Wolframin might regulate MCT1/Basigin protein processing and/or stability also outside the retina causing other symptoms associated with Wolfram syndrome. In fact, MCT1 is strongly enriched in brain astrocytes and oligodendrocytes and sustain the lactate shuttle towards neuronal cells and their metabolic fitness and survival (Fünfschilling et al., 2012; Lee et al., 2012). Thus, MCT1 loss in the CNS might contribute to diffuse neurodegeneration underlying the progressive brain volume loss occurring in WS1 patients. Finally, MTC1 is the most abundantly expressed lactate transporter in peripheral nerves and its inactivation in Schwann cells leads to hypomyelination and functional deficits in sensory, but not motor, peripheral nerves (Jha et al., 2020). Intriguingly, WS1 patients might develop peripheral neuropathy with increased impairment of the sensory component (Liu et al., 2006). Future studies are warranted to determine lactate metabolism dysregulation in brain and peripheral neurons in *Wfs1* mutant mice.

These results have also profound implications on future strategies for gene therapy in WS1. In fact, given the selective RGC loss, research activities are in progress to establish gene therapy approaches to express a functional *Wfs1* gene copy in RGCs in *Wfs1* mutant mice. However, given our results we expect that this design will not efficiently prevent RGC dysfunction and subsequent loss. In contrast, *Wfs1* gene activity should be restored in glial cells of the retina and optic nerve in order to correct the trafficking of energy metabolites and sustain the bioenergetics of RGC cell bodies and axons. Altogether, by revealing new pathophysiological mechanisms underlying visual loss in *Wfs1* mutant mice these findings uncover unprecedented therapeutic directions for WS1 calling for the exploitation of pharmacological strategies for metabolic support based on boosting glucose alternative energy substrates (Camandola and Mattson, 2017; Beard et al., 2022).

## Acknowledgments

We thank A. Plaas and E. Vasar for providing us the *Wfs1* mutant mice, P. Podini for expert support on EM imaging and I. Viganò and G. Zerbini for OCT and FA analyses. We are thankful to G. Frontino, L. Piemonti, P. Barboni and M.L. Cascavilla for helpful discussion. We acknowledge D. Bonanomi and members of the Broccoli’s lab for generous support and advice. This work was supported by a private family donation financing the work on Wolfram syndrome in V.B. lab at OSR and PON ELIXIR CNR-BiOMICS (PIR01_00017), Elixir Implementation Study Proteomics 2021-23 and Italian Ministry of Health (RF2019-12370396) to P.M.

## Author contributions

G.R. performed the experiments and analyzed the data with the help of S.M. and N.N.V. G.O. analyzed protein levels by immunoblot in retinal lysates; A.I. executed the biochemical assays and analyzed the results. V.C. performed the electrophysiological recordings and analyzed the data together with L.L. E.B. carried out the RNA-seq computational analysis and data mining. D.D.S. and L.B. carried out the label-free proteomics under the supervision of P.L.M. who elaborated the results. S.G.G. designed and finalized the molecular cloning. V.B. designed the study, supervised research, and acquired funding. V.B. wrote the manuscript with input from all co-authors. All authors discussed the data and manuscript.

## Declaration of Interests

The authors declare no competing interests.

## METHODS

### Animals

The *Wfs1*^exon8del^ targeted mouse model was a kind gift of M. Plaas and colleagues (Luuk et al., 2008). Mice were genotyped by multiplex PCR for assessing both WT and mutant alleles using primers WfsKO_wtF2 5’ TTGGCTTGTATTTGTCGGCC, NeoR1 5’ GACCGCTATCAGGACATAGCG and WfsKO_uniR2 5’ CCCATCCTGCTCTCTGAACC (Luuk *et al*, 2009). Mice were maintained on a C57BL/6J background at San Raffaele Scientific Institute Institutional mouse facility (Milan, Italy). All procedures were performed according to protocols approved by the internal IACUC and reported to the Italian Ministry of Health according to the European Communities Council Directive 2010/63/EU.

### Optical Coherence Tomography (OCT) analysis and Fluorescein Angiography (FA)

Optical coherence tomography (OCT) and fluorescein angiography (FA) were performed in collaboration with the CIS (Experimental Clinical Imaging) facility of San Raffaele Scientific Institute (Milan) as previously described (Buccarello et al., 2017), using the Micron IV instrument (Phoenix Research Laboratories, Pleasanton, CA, United States). Briefly, after anaesthesia, mydriasis was induced by administering a drop of tropicamide 0.5% (Visumidriatic, Tibilux Pharma, Milan, Italy) in each eye. OCT images were acquired by performing a circular scan of 550 μm of diameter around the optic nerve head. Both eyes were examined, and the results were averaged. The segmentation of retinal layers was performed using Insight software (Phoenix Research Laboratories, Pleasanton, CA, United States), OCT was followed by the FA study. A solution of 1% fluorescein (5 ml/kg Monico S.p.A., Venezia, Italy) was administered by a single intraperitoneal injection (100 μL). For each animal, the images of central and peripheral retinal vasculature were acquired.

### Photopic electroretinogram (pERG) recordings

pERGs were recorded after 10 min of light adaptation under intraperitoneal anaesthesia (80 mg/kg ketamine, 10 mg/kg xylazine). Pupils were dilated with 0.5% tropicamide and moisturized with ophthalmic gel (2% hydroxypropylmethylcellulose) to avoid eye drying. Body temperature was maintained with a homeothermic blanket system at 36.5 ± 0.5 °C. pERG was recorded from one eye at a time using a corneal electrode connected via flexible cables to a Micromed amplifier, as reported previously (Marenna et al., 2020). Each session included 3 trains of 10 flash stimuli (with 130 mJ intensity, 10 ms duration and 0.5 Hz frequency) delivered with a flash photostimulator (Micromed, Mogliano Veneto, Italy) placed at 15 cm from the stimulated eye. pERGs were acquired with Micromed System Plus Evolution at a sampling frequency of 4096 Hz, coded with 16 bits, bandpass-filtered (5–100 Hz), and notch filtered (50 Hz). Implicit time of PhNR was measured, together with its corresponding amplitude (from baseline to negative PhNR peak). Left and right eyes were averaged to obtain single values for each animal.

### Visual Evoked Potential (PEV) recordings

Before recording procedures, mice were placed in a dark room and allowed to adapt to darkness for 5 minutes. Mice were intraperitoneally anesthetized (80 mg/kg ketamine, 10 mg/kg xylazine) and adequate level of anesthesia was verified by checking for the presence of tail-pinching reflex. Body temperature was maintained at 36.5 ± 0.5 °C by a homeothermic blanket system with a rectal thermometer probe. Both eyes were dilated with 0.5% tropicamide and protected using ophthalmic gel. Non-invasive epidermal VEPs were recorded using a 6 mm Ø Ag/AgCl cup electrode placed on the shaved scalp over V1, contralateral to the stimulated eye (1 mm anterior to interaural line and 2.5 mm contralateral to flash stimulation) and a needle electrode was inserted in the snout for reference. The cup was fixed with electro-conductive adhesive paste and at the end of first eye recording, it was placed on the opposite hemisphere to acquire VEPs from the contralateral eye, as described previously (Marenna et al, 2019). For each VEP recording session, 3 trains of 20 flash stimuli (with 260 mJ intensity, 10 ms duration and 1 Hz frequency) were delivered with a flash photostimulator (Micromed, Mogliano Veneto, Italy) placed at 15 cm from the stimulated eye, while the contralateral eye was covered. VEPs from both eyes were acquired and measured offline with Micromed System Plus Evolution software at a sampling frequency of 4096 Hz, coded with 16 bits, bandpass filtered (5–100 Hz) and notch filtered (50 Hz). Latency of the first negative peak (N1) and peak-to-peak amplitude (N1-P2) were measured, and then left and right eyes were averaged to obtain single values for each mouse.

### Visual acuity assessment using the optomotor reflex system

The spatial frequency threshold (“visual acuity”) of the optomotor reflex index was determined using the Optodrum instrumentation (Striatech, Germany). Briefly, freely moving mice were placed on an elevated platform and exposed to vertical sine-wave gratings rotating at 12°/s. Spatial frequency of the grating at full contrast was gradually increased until the mice no longer were tracking the grating with reflexive head movements in concert with the rotation. The results are averaged for right- and left-eye acuity. Mice were tested within 5 hours of their daylight hour onset.

### Transmission electron microscopy analysis and morphometric evaluation of optic nerve fibers

The samples were fixed for 1 h at 4 °C with 2,5% glutaraldehyde, 4% PFA in 0.1 M cacodylate buffer, pH 7.35. Fixation buffer was removed and cells washed 3x with 0.1 M cacodylate buffer, pH 7.35 post-fixed in 2% OsO4 solution for 1 h, and stained in 1% uranyl acetate for 1 h at room temperature. After dehydration, specimens were embedded in Epon resin. Ultra-thin sections (about 70 nm) were stained with uranyl acetate and lead citrate and were examined by electron microscopy (Fei Talos L120CG2). For fiber counts, regions were randomly selected to minimize the bias due to fibre size distribution along the nerve. Over 200 nerve fibres (on average) were counted from individual mice (4–6 animals from each group). All nerve fibres and axon analysis were executed using AxonJ plugin of Fiji software (Marenna et al., 2019). Total Axon area quantifications were performed on binned and gaussian blurred images and the total was normalized per field of view area. Area based g-ratio (axon area divided by axon area plus myelin area) were used.

### Tissue collection and immunohistochemistry

Animals were euthanized and the eyes and brains collected and subjected to the different procedures based on the type of analysis. In particular, after visual functional and structural analysis, mice were sacrificed and eyes with optic nerves were harvested for light and electron microscopic evaluation. Dissected retinas were fixed in 4% (w/v) PFA overnight, washed in PBS and soaked in cryoprotective solution (30% sucrose in PBS) overnight. Then, embedding in OCT Compound (VWR) was performed and eye tissues were cryosectioned in 18um-thick sections and subjected to immunostaining. Where indicated, RGC density in the ganglion cell layer was determined for each eye by counting the number of cells over a 300 μm distance in serial sections representative of the full retina. For protein/RNA isolation, freshly collected retinas were flash frozen on dry-ice and stored until extraction. For all other tissues collection mice were previously anesthetized and transcardially perfused with 0.1 M phosphate buffer (PB) at room temperature (RT) at pH 7.4. Subsequently, tissues were collected and post-fixed (for IF/IHC analysis) or flash frozen on dry ice (for RNA/Protein extraction) as needed. For immunohistochemistry, retinas were fixed in 4% of PFA, soaked in cryoprotective solution (30% sucrose in PBS), flash frozen and cut into 50 μm slices. Next, the slices were permeabilized with 3% H2O2, 10% methanol and 2% of triton for 20 minutes. After 3 washes the slices were treated with the blocking solution (3% BSA, 0,1% tween 20) and incubated with the primary antibody over night at 4°C. For the immunofluorescence the secondary antibody was added, and the slices mounted for the imaging.

### Western blot analysis

Samples were homogenized in RIPA Buffer (Tris-HCl 100mM pH 7.4, NaCl 150mM, EGTA 1mM, Triton 0.5%, SDS 0.1% with protease (2%) and phosphatase (10%) inhibitors (Roche)). After a 30 min incubation at 4°C, lysates were centrifuged at 13.000g for another 30 min and the supernatant was collected into new Eppendorf tubes. Present proteins were then quantified using the Pierce BCA Protein Assay (Thermo Scientific). Electrophoretic run of samples containing 30μg of proteins was performed on precast NuPAGE 3-8% or 4-12% Tris-Acetate Protein Gels (Thermo Fisher Scientific), or on home-made acrylamide gels 8-15%. After the run (30-60 min at 150V), proteins were transferred to a nitrocellulose membrane using the Trans-Blot Turbo RTA Mini Nitrocellulose Transfer Kit (Biorad). To block the unspecific sites, membranes were incubated for 1h at RT in 5% non-fat dry milk (5% BSA where indicated), 0,1% PBS-Tween solution and then overnight at 4°C in primary antibody (antibodies specifics are listed in **Table 2**), diluted in the same solution. The day after, membranes were incubated for 1h at RT with the secondary antibody (Polyclonal Goat Anti-Rabbit Immunoglobulins/HRP, Dako; Polyclonal Goat Anti-Mouse Immunoglobulins/HRP, Dako) prepared in the same blocking solution at a dilution of 1:10.000. After several washes, membranes were revealed with SuperSignal West Pico PLUS Chemiluminescent Substrate ECL (Thermo Fisher Scientific). Chemiluminescence was detected using ChemiDoc Gel Imaging System (Biorad). Finally, band densitometry relative to control was calculated using ImageLab software, normalized on housekeeping as indicated in each figure (CALNEXIN and ACTIN).

### Co-immunoprecipitation assays

For the Co-IP, HEK-293T cells were cultured in 10cm dishes. Expression vectors with *WFS*-mCherry and *MCT1*-V5 were used for transient transfection in cells. Cells were lysed in IP buffer (Tris-HCl pH 7.5 20mM, NaCl 150mM, EDTA 1mM, EGTA 1mM, 1% Triton X-100, Protease inhibitors and Phosphatase inhibitors in milli-Q) for 30 minutes at 4°C, vortexing every 10 minutes. Then the lysed cells were centrifuged for 10 minutes at 14.000 x g at 4°C to pellet the debris. The supernatant was collected, and proteins were quantified using the Pierce BCA Protein Assay kit (ThermoFisher Scientific). From each sample, 60μg were kept as the INPUT, which corresponds to the cell lysate before being co-immunoprecipitated. After that, the same amount of the remaining protein was taken from all samples (accordingly to that of the limiting sample) and brought to a final volume of 450μL with IP buffer. Generally, not less than 2mg were used as starting material for the subsequent IP procedure. Protein lysates were pre-cleared 1h at 4°C on a rotating wheel with 50μl of previously washed Dynabeads Protein G (Invitrogen, Thermo Fisher Scientific), to avoid aspecific bindings. The samples were then centrifuged at max speed for 10 minutes to precipitate the beads. The supernatant was then collected and incubated overnight at 4°C on the rotating wheel with the antibodies anti-V5 or anti-RFP (the antibodies with their relative dilutions are listed in **Table S2**). The day after, each sample was incubated with 100μl of washed beads for 2h at 4°C on the rotating wheel, in order to precipitate the proteins bound to the antibodies. After that, using the magnetic particle concentrator (DynaMag-Spin, Invitrogen), the fluid was collected as the unbound, representing the protein fraction that was not co-immunoprecipitated. Then, still using the magnet, the beads were washed 8 times with 1mL of IP buffer. The beads were resuspended in 100μl of Laemmli buffer 1X and boiled at 95°C for 5 minutes, to detach the proteins from the beads. The co-immunoprecipitated fraction was separated from the beads with the magnet and collected in a new Eppendorf. The unbounds were supplemented with 100μl of Laemmli buffer 4X, while the INPUTs were additioned with Laemmli buffer 4X and an appropriate amount of H2O to bring the concentration of the buffer to 1X, reaching a final volume of 32μl. The negative controls were obtained by immunoprecipitating the double transfected cells lysates with an isotype control (a non-specific antibody of the same species and isotype of the specific antibody used for the Co-IP).

### RNA-Seq and computational analysis

For transcriptome analysis, total RNA was extracted and purified with the QIAGEN RNeasy Micro Kit from freshly enucleated eyes from 8-month-old mice (n=3 control vs n=3 *Wfs1*-deficient mice). Sequencing was performed by GENEWIZ sequencing company (Germany). FASTQ reads were quality checked with FastQC (v 0.11.9) and adaptors trimmed with Trimmomatic (v 0.40) (Bolger et al., 2014), specifying default parameters. High-quality trimmed reads were mapped to the mm10 reference genome with STAR aligner (v 2.7.9.a) using the latest GENCODE main annotation file. The construction of the gene count matrix was done using featureCounts (Liao et al., 2014) on .bam files generated by STAR, counting the reads associated to ’exons’ features per gene. Differential gene expression on the normalized gene count matrix by RPKM method was performed with DESeq2 (v 3.13). DEGS (differentially expressed genes) were extracted by filtering on Pvalue adjusted (using 0.05 as threshold value), corrected by the Benjamini Hochberg test. GO enrichment analysis on DEGS was performed with Gprofiler2 (v 0.2.0) (Love *et* al., 2014); only overrepresented categories with corrected Pvalue were kept, using the previously described method. Downstream statistics and Plotting were performed within the R (v 4.0.1) environment. Heatmaps were generated with ComplexHeatmap (v 3.0.215).

### qRT-PCR analysis

RNA was extracted using the TRI Reagent isolation system (Sigma-Aldrich) according to the manufacturer’s instructions. For quantitative RT-PCR (qRT-PCR), one microgram of RNA was reverse transcribed using the ImProm-II Reverse Transcription System (Promega), thereafter qRT-PCR was performed in triplicate with custom designed oligos (**Table S3**) using the CFX96 Real-Time PCR Detection System (Bio-Rad, USA). using the Titan HotTaq EvaGreen qPCR Mix (BIOATLAS). Obtained cDNA was diluted 1:10 and was amplified in a 16μl reaction mixture containing 2 μl of diluted cDNA, 1× Titan Hot Taq EvaGreen qPCR Mix (Bioatlas, Estonia) and 0.4 mM of each primer. Analysis of relative expression was performed using the ΔΔCt method, using 18S rRNA as housekeeping gene and CFX Manager software (Bio-Rad, USA).

### Proteomics sample preparation and Liquid chromatography-tandem mass spectrometry (LC-MS/MS)

Wild-type (WT) and *Wfs1* mutant retinal mouse samples sampling at 8 months were analysed in label-free proteomics approach. A total of 6 samples were prepared for proteomics analyses by means EasyPep^TM^ Mini MS Sample Prep Kit (Thermo Fisher Scientific, Rock ford, IL, USA) for proteins extraction, reduction/alkylation and digestion by trypsin and Lys-C combination enzymes. The protein concentration was measured by Pierce BCA Protein Assay kit (Thermo Fisher Scientific, Rock ford, IL, USA). The peptides mixtures were clean-up to obtain detergent-free samples with the same kit, and then resuspended in 0.1% formic acid (Sigma-Aldrich Inc., St. Louis, MO, USA) for mass spectrometry analysis. The peptide samples were analysed on an LTQ-OrbitrapXL mass spectrometer Kit (Thermo Fisher Scientific, Rock ford, IL, USA) coupled to the Eksigent nanoLC-Ultra® 2D system (AB Sciex, Dublin, CA, USA). Briefly, 0.8 µg peptides were loaded on a trap (200 x 500 µm ChromXP C18-CL, 3 µm, 120 Å) and eluted on a reversed-phase column (75 µm x 15 cm ChromXP C18-CL, 3 µm, 120 Å) using a acenotrile gradien;B (Eluent A: 0.1% formic acid in water; Eluent B: 0.1% formic acid in acetonitrile): 5-10% in 3 min, 10-40% in 87 min, 40-95% in 10 min, and holding at 95% B for 8 min; flow rate was 300nL/minute. The spray capillary voltage was set at 1.7 kV, and the ion transfer capillary temperature was held at 220 °C. Full MS spectra were recorded over 400-1600 m/z range in positive ion mode, resolving power of 30,000, followed by five tandem mass spectrometry (MS/MS) on the top most intense ions selected from the full MS spectrum, using a dynamic exclusion for MS/MS analysis.

### LC-MS/MS data handling and data processing

Raw files generated were processed by Proteome Discoverer 2.1 (Thermo Fisher Scientific) using Sequest HT search engine. The experimental MS/MS spectra were compared with the theoretical mass spectra obtained by *in silico* digestion of a *Mus musculus* protein database containing 55282 sequences (April 2022 UniProt version; www.uniprot.org). Searching criteria were used: Trypsin and Lys-C enzyme, two missing cleavage, precursor mass tolerance ± 50 ppm, fragment mass tolerance ± 0.8 Da. The Percolator node with a target-decoy strategy was selected to give a false discovery rate of ≤ 0.01 (strict) The Peptide Spectrum Matches (PSM) values of identified proteins were used for label-free quantitation approach. The whole matrix was reduced by Linear Discriminant Analysis (LDA) and a pairwise comparison (3 WT *vs* 3 *Wfs1*-KO) was performed and extracted descriptors (p-value < 0.05). PSM. Differential expressed proteins were obtained by MAProma Algorithms (Sereni et al., 2019) and filtered by p-value (less than 0.05).

### Lactate and ATP biochemical assays

Lactate level in retinae and optic nerve was measured by the Lactate-Glo^TM^ Assay (Promega). Samples were disaggregated for 20–30 seconds using a tissue tearor homogenizer in 50mM Tris, pH 7.5 pre-mixed with inactivation Solution (0.6N HCl). Then the samples were neutralized by ml neutralization solution (1M Tris base). After, 50 µl of prepared samples was added to 50 µl of working solution before reading the plate on the luminometer Mithras LB 940 (Berthold Technologies, Switzerland).

ATP concentration was determined using ATP Assay Kit (Abcam). Samples were homogenized in 100 μL of ATP Assay Buffer with a tissue tearor. Then, the samples were centrifuged for 2-5 minutes at 4°C at 13,000 g using a cold microcentrifuge to remove any insoluble material and the supernatant was collected in a new tube. After 50 µl of prepared samples was added to 50 µl of working solution before reading the optical density (OD) at OD 570 nm using the Epoch Microplate Spectrophotometer (BioTek).

### Statistics

All values are expressed as mean ± standard error (SEM) of at least 3 independent experiments, as indicated. All the statistical analysis was carried out with Prism 8 software (GraphPad software). Differences between means were analyzed using the Student’s *t*-test for experimental designs with less than three groups and one variable, alternatively we applied one-way/two-way ANOVA followed by Bonferroni and Tukey post hoc test. Electrophysiological comparisons between WT and KO mice were performed using Student’s t test for homoscedastic samples or Welch’s t-test for heteroscedastic samples, after testing for the equality of variances with Levene’s test. The null hypothesis was rejected when p < 0.05. In the graphs “ns” indicates non-significant differences, * indicates significant differences with *p* < 0.05, ** indicates significant differences with *p* < 0.01, *** indicates significant differences with *p* < 0.001. In case of unequal variances (heteroscedasticity), Welch test was applied.

---

**Supplementary Figure 1:**
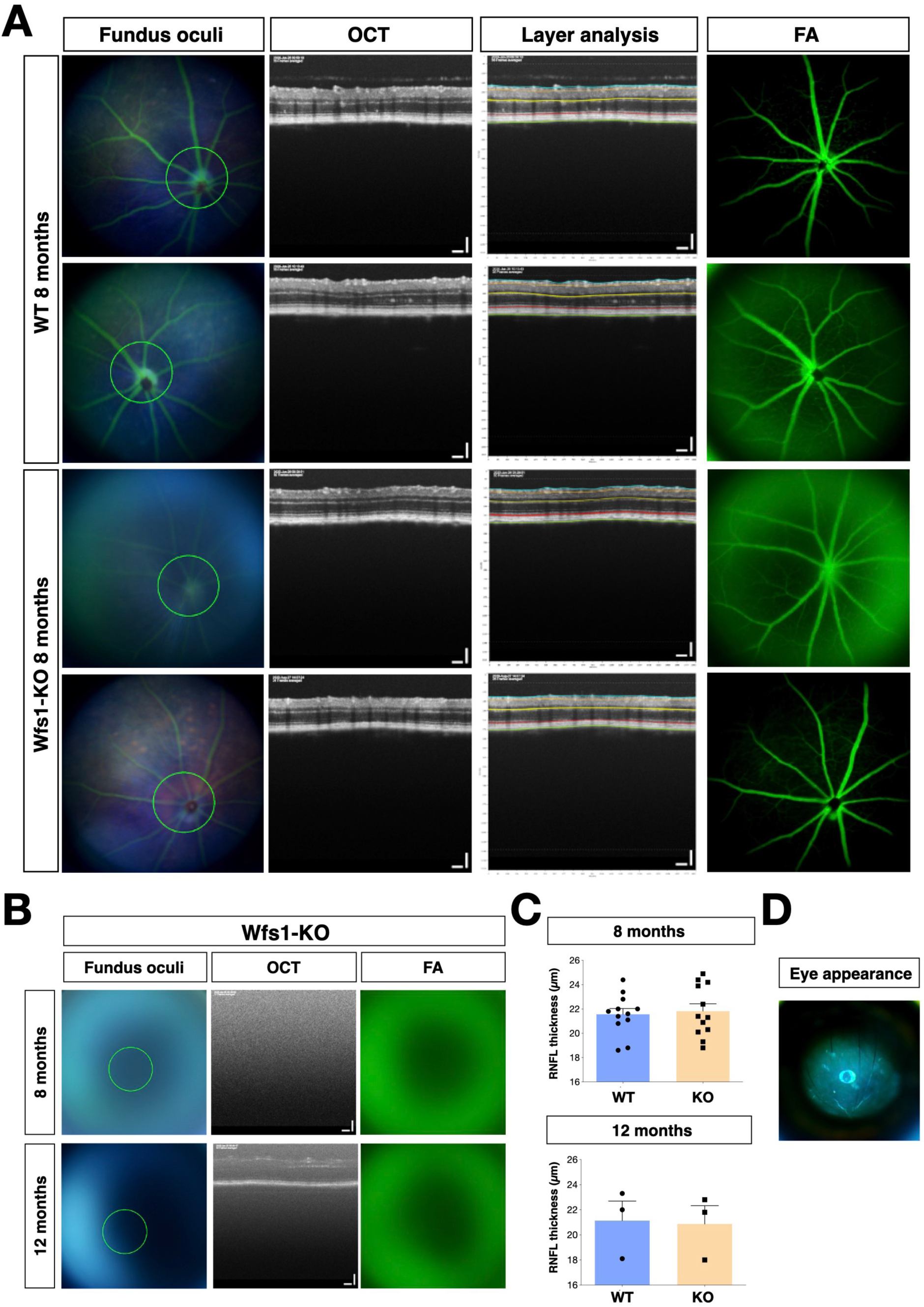
Characterization of eye morphology in *Wfs1* mutant mice. (A,B) Representative images from fundus oculi, OCT, layer analysis and fluorescein angiography (FA) in wild-type (WT) and *Wfs1* mutant mice at 8 months of age. (C) Quantification of RNFL thickness expressed in μm in 8 month (n = 12, *p* = 0,73, unpaired *t*-test, top) or 12 month (n = 3, *p* = 0,90, unpaired *t*-test, bottom) old animals. (D) Representative image of *Wfs1* mutant mouse eye with diffuse cataract and corneal haze.

**Supplementary Figure 2:**
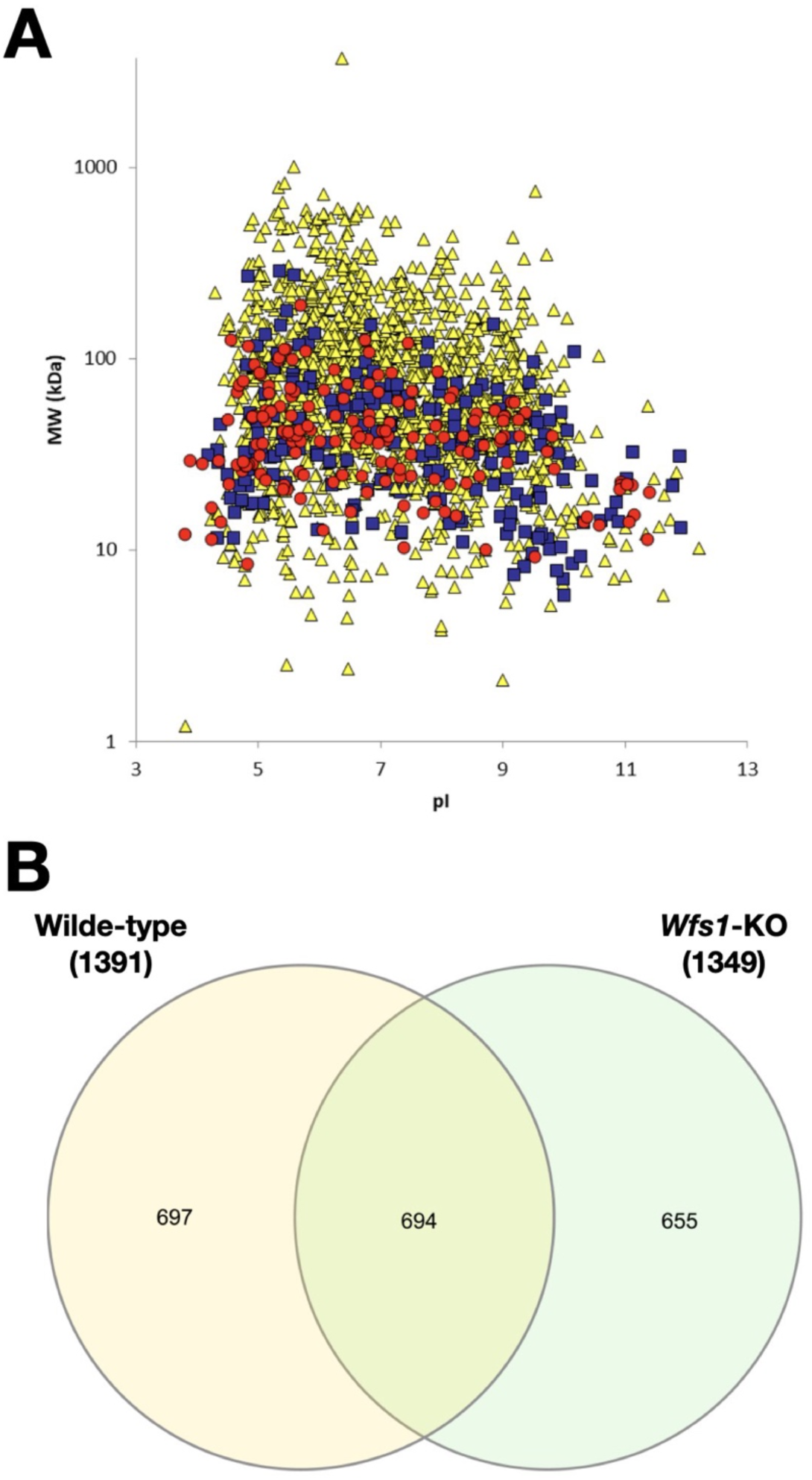
General analysis of the proteomics output and identified proteins. (A) Virtual 2D maps, based on theroretical MW and pI of proteins. Color/shape code is assigned to each protein according to the corresponding PSM value; PSM=1, >1 and ≤5, and >5 are depicted in yellow/triangle, blue/square, and red/circles, respectively. B) Venn diagram of identified protein in wild-type (WT) and *Wfs1* mutant (KO) retinal samples. The area of intersection is related to shared proteins.

## Notes

### Competing Interest Statement

The authors have declared no competing interest.

